# MINT: A multivariate integrative method to identify reproducible molecular signatures across independent experiments and platforms

**DOI:** 10.1101/070813

**Authors:** F. Rohart, A. Eslami, N. Matigian, S. Bougeard, K-A. Lê Cao

## Abstract

**Background:** Molecular signatures identified from high-throughput transcriptomic studies often have poor reliability and fail to reproduce across studies. One solution is to combine independent studies into a single integrative analysis, additionally increasing sample size. However, the different protocols and technological platforms across transcriptomic studies produce unwanted systematic variation that strongly confounds the integrative analysis results. When studies aim to discriminate an outcome of interest, the common approach is a sequential two-step procedure; unwanted systematic variation removal techniques are applied prior to classification methods.

**Results:** To limit the risk of overfitting and over-optimistic results of a two-step procedure, we developed a novel multivariate integration method, *MINT*, that simultaneously accounts for unwanted systematic variation and identifies predictive gene signatures with greater reproducibility and accuracy. In two biological examples on the classification of three human cell types and four subtypes of breast cancer, we combined high-dimensional microarray and RNA-seq data sets and MINT identified highly reproducible and relevant gene signatures predictive of a given phenotype. MINT led to superior classification and prediction accuracy compared to the existing sequential two-step procedures.

**Conclusions:** *MINT* is a powerful approach and the first of its kind to solve the integrative classification framework in a single step by combining multiple independent studies. *MINT* is computationally fast as part of the mixOmics R CRAN package, available at http://www.mixOmics.org/mixMINT/ and http://cran.r-project.org/web/packages/mixOmics/.

## 1 Introduction

High-throughput technologies, based on microarray and RNA-sequencing, are now being used to identify biomarkers or gene signatures that distinguish disease subgroups, predict cell phenotypes or classify responses to therapeutic drugs. However, few of these findings are reproduced when assessed in subsequent studies and even fewer lead to clinical applications (Pihur *et al.*, 2008; Kim *et al.*, 2016). The poor reproducibility of identified gene signatures is most likely a consequence of high-dimensional data, in which the number of genes or transcripts being analysed is very high (often several thousands) relative to a comparatively small sample size being used (< 20).

One way to increase sample size is to combine raw data from independent experiments in an integrative analysis. This would improve both the statistical power of the analysis and the reproducibility of the gene signatures that are identified (Lazar *et al.*, 2012). However, integrating transcriptomic studies with the aim of classifying biological samples based on an outcome of interest (integrative classification) has a number of challenges. Transcriptomic studies often differ from each other in a number of ways, such as in their experimental protocols or in the technological platform used. These differences can lead to so-called ‘batch-effects’, or systematic variation across studies, which is an important source of confounding (Gagnon-Bartsch and Speed, 2012). Technological platform, in particular, has been shown to be an important confounder that affects the reproducibility of transcriptomic studies (Shi *et al.*, 2006). In the MicroArray Quality Control (MAQC) project, poor overlap of differentially expressed genes was observed across different microarray platforms (~ 60%), with low concordance observed between microarray and RNA-seq technologies specifically (Su *et al.*, 2014). Therefore, these confounding factors and sources of systematic variation must be accounted for, when combining independent studies, to enable genuine biological variation to be identified.

The common approach to integrative classification is sequential. A first step consists of removing batch-effect by applying for instance ComBat (Johnson *et al.*, 2007), FAbatch (Hornung *et al.*, 2016), Batch Mean-Centering (Sims *et al.*, 2008), LMM-EH-PS (Listgarten *et al.*, 2010), RUV-2 (Gagnon-Bartsch and Speed, 2012) or YuGene (Lê Cao *et al.*, 2014). A second step fits a statistical model to classify biological samples and predict the class membership of new samples. A range of classification methods also exists for these purposes, including machine learning approaches (e.g. random forests, Breiman 2001 or Support Vector Machine) as well as multivariate linear approaches (Linear Discriminant Analysis LDA, Partial Least Square Discriminant Analysis PLSDA, Barker and Rayens 2003, or sparse PLSDA (Lê Cao *et al.*, 2011)).

The major pitfall of the sequential approach is a risk of over-optimistic results from overfitting of the training set. This leads to signatures that cannot be reproduced on test sets. Moreover, most proposed classification models have not been objectively validated on an external and independent test set. Thus, spurious conclusions can be generated when using these methods, leading to limited potential for translating results into reliable clinical tools (Kim *et al.*, 2016). For instance, most classification methods require the choice of a parameter (e.g. sparsity), which is usually optimised with cross-validation (data are divided into k subsets or ‘folds’ and each fold is used once as an internal test set). Unless the removal of batch-effects is performed independently on each fold, the folds are not independent and this leads to over-optimistic classification accuracy on the internal test sets. Hence, batch removal methods must be used with caution. For instance, ComBat can not remove unwanted variation in an independent test set alone as it requires the test set to be normalised with the learning set in a transductive rather than inductive approach (Hughey and Butte, 2015). This is a clear example where over-fitting and over-optimistic results can be an issue, even when a test set is considered.

To address existing limitations of current data integration approaches and the poor reproducibility of results, we propose a novel Multivariate INTegrative method, *MINT*. *MINT* is the first approach of its kind that integrates independent data sets while *simultaneously*, accounting for unwanted (study) variation, classifying samples and identifying key discriminant variables. *MINT* predicts the class of new samples from external studies, which enables a direct assessment of its performance. It also provides insightful graphical outputs to improve interpretation and inspect each study during the integration process.

We validated MINT in a subset of the MAQC project, which was carefully designed to enable assessment of unwanted systematic variation. We then combined microarray and RNA-seq experiments to classify samples from three human cell types (human Fibroblasts (Fib), human Embryonic Stem Cells (hESC) and human induced Pluripotent Stem Cells (hiPSC)) and from four classes of breast cancer (subtype *Basal, HER2, Luminal A* and *Luminal B* (Parker *et al.*, 2009)). We use these datasets to demonstrate the reproducibility of gene signatures identified by *MINT*.

## 2 Results

### 2.1 Validation of the *MINT* approach to identify signatures agnostic to batch effect

The MAQC project processed technical replicates of four well-characterised biological samples A, B, C and D across three platforms. Thus, we assumed that genes that are differentially expressed (DEG) in every single platform are true positive. We primarily focused on identifying biomarkers that discriminate C vs. D, and report the results of A vs. B in the Supplemental Material S4.1. Differential expression analysis of C vs. D was conducted on each of the three microarray platforms using ANOVA, showing an overlap of 1,385 DEG (FDR < 10^−3^, Benjamini and Hochberg 1995), which we considered as true positive. This corresponded to 62.6% of all DEG for Illumina, 30.5% for AffyHuGene and 21.0% for AffyPrime (Figure S3). We observed that conducting a differential analysis on the concatenated data from the three microarray platforms without accommodating for batch effects resulted in 691 DEG, of which only 56% (387) were true positive genes. This implies that the remaining 44% (304) of these genes were false positive, and hence were not DE in at least one study. The high percentage of false positive was explained by a Principal Component Analysis (PCA) sample plot that showed samples clustering by platforms (Figure S3), which confirmed that the major source of variation in the combined data was attributed to platforms rather than cell types.

MINT selected a single gene, BCAS1, to discriminate the two biological classes C and D. BCAS1 was a true positive gene, as part of the common DEG, and was ranked 1 for Illumina, 158 for AffyPrime and 1,182 for AffyHuGene. Since the biological samples C and D are very different, the selection of one single gene by MINT was not surprising. To further investigate the performance of MINT, we expanded the number of genes selected by MINT, by decreasing its sparsity parameter (see Methods), and compared the overlap between this larger MINT signature and the true positive genes. We observed an overlap of 100% for a MINT signature of size 100, and an overlap of 89% for a signature of size 1, 385, which is the number of common DEG identified previously. The high percentage of true positive selected by MINT demonstrates its ability to identify a signature agnostic to batch effect.

### 2.2 Limitations of common meta-analysis and integrative approaches

**Figure 1:**
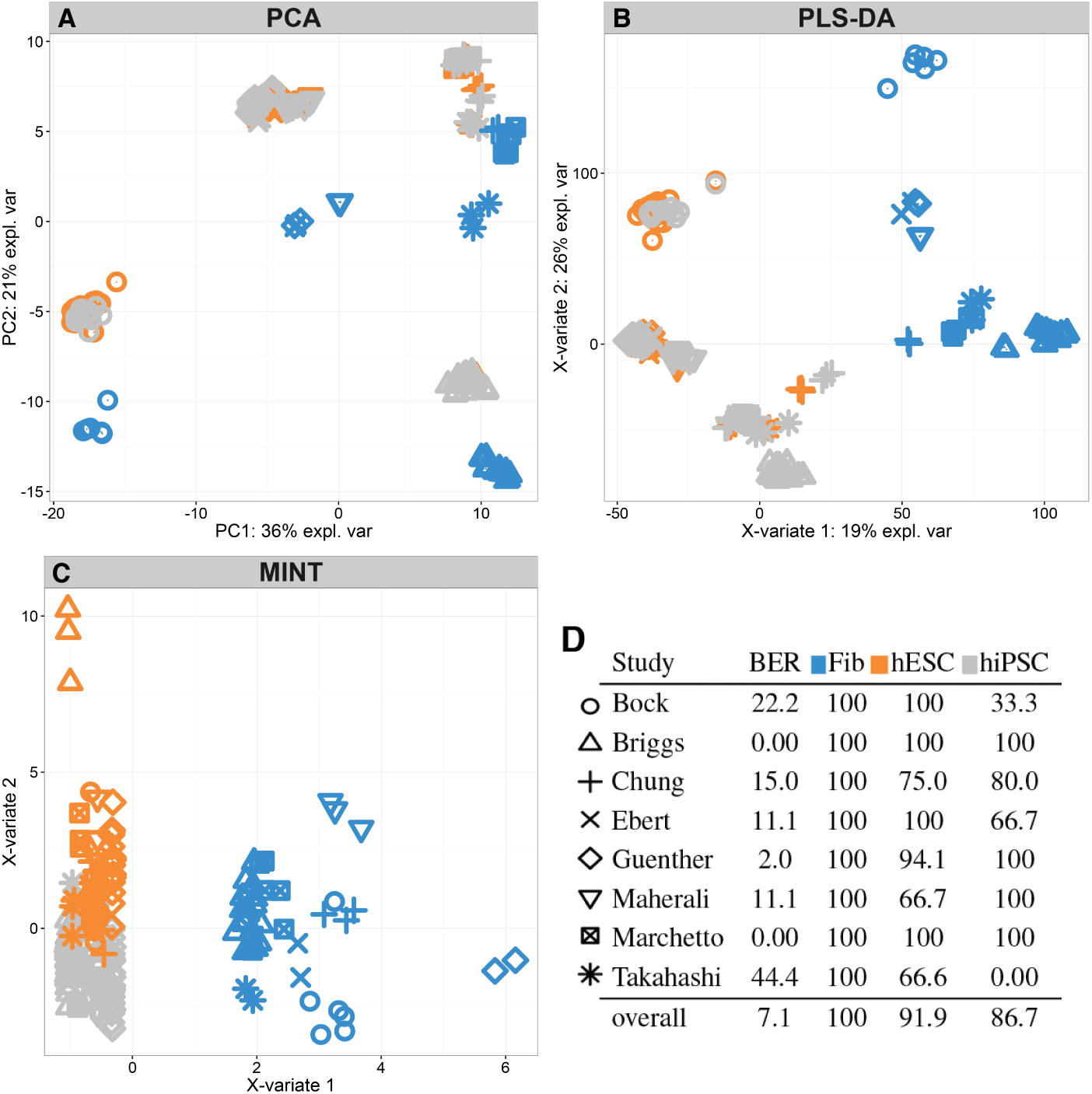
Stem cell study. (A) PCA on the concatenated data: a greater study variation than a cell type variation is observed. (B) PLSDA on the concatenated data clustered Fibroblasts only. (C) *MINT* sample plot shows that each cell type is well clustered, (D) *MINT* performance: BER and classification accuracy for each cell type and each study.

A meta-analysis of eight stem cell studies, each including three cell types (Table 1, stem cell training set), highlighted a small overlap of DEG lists obtained from the analysis of each separate study (FDR < 10^*−*5^, ANOVA, Supplemental Information S5.2). Indeed, the Takahashi study with only 24 DEG limited the overlap between all eight studies to only 5 DEG. This represents a major limitation of merging pre-analysed gene lists as the concordance between DEG lists decreases when the number of studies increases.

One alternative to meta-analysis is to perform an integrative analysis by concatenating all eight studies. Similarly to the MAQC analysis, we first observed that the major source of variation in the combined data was attributed to study rather than cell type (Figure 1A). PLS-DA was applied to discriminate the samples according to their cell types, and it showed a strong study variation (Figure 1B), despite being a supervised analysis. Compared to unsupervised PCA (Figure 1A), the study effect was reduced for the fibroblast cells, but was still present for the similar cell types hESC and hiPSC. We reached similar conclusions when analysing the breast cancer data (Supplemental Information S6.2).

### 2.3 *MINT* outperforms state-of-the-art methods

We compared the classification accuracy of *MINT* to sequential methods where batch removal methods were applied prior to classification methods. In both stem cell and breast cancer studies, MINT led to the best accuracy on the training set and the best reproducibility of the classification model on the test set (lowest Balanced Error Rate, BER, Figure 2, S12). In addition, MINT consistently ranked first as the best performing method, followed by ComBat+sPLSDA with an average rank of 4.5 (Figure S4).

**Figure 2:**
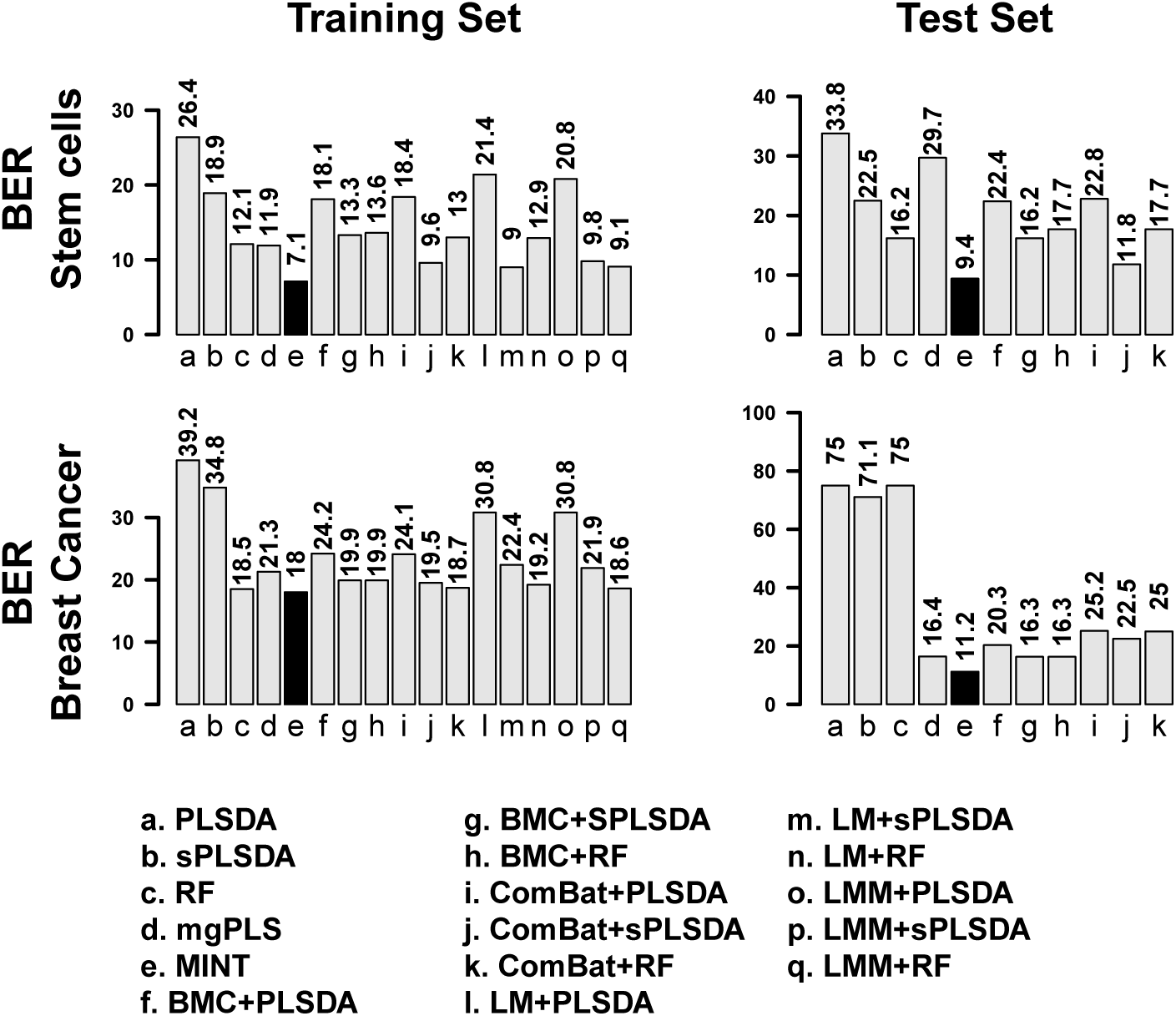
Classification accuracy for both training and test set for the stem cells and breast cancer studies (excluding PAM50 genes). The classification Balanced Error Rates (BER) are reported for all sixteen methods compared with MINT (in black).

On the stem cell data, we found that fibroblasts were the easiest to classify for all methods, including those that do not accommodate unwanted variation (PLS-DA, sPLS-DA and RF, Figure S6). Classifying hiPSC vs. hESC proved more challenging for all methods, leading to a substantially lower classification accuracy than fibroblasts.

The analysis of the breast cancer data (excluding PAM50 genes) showed that methods that do not accommodate unwanted variation were able to rightly classify most of the samples from the training set, but failed at classifying any of the four subtypes on the external test set. As a consequence, all samples were predicted as *LumB* with PLS-DA and sPLS-DA, or *Basal* with RF (Figure S12). Thus, RF gave a satisfactory performance on the training set (BER = 18.5), but a poor performance on the test set (BER= 75).

Additionally, we observed that the biomarker selection process substantially improved classification accuracy. On the stem cell data, LM+sPLSDA and *MINT* outperformed their non sparse counterparts LM+PLSDA and mgPLS (Figure 2, BER of 9.8 and 7.1 vs. 20.8 and 11.9), respectively.

Finally, *MINT* was largely superior in terms of computational efficiency. The training step on the stem cell data which includes 210 samples and 13, 313 was run in 1 second, compared to 8 seconds with the second best performing method ComBat+sPLS-DA (2013 MacNook Pro 2.6Ghz, 16Gb memory). The popular method ComBat took 7.1*s* to run, and sPLS-DA 0.9*s*. The training step on the breast cancer data that includes 2, 817 samples and 15, 755 genes was run in 37*s* for MINT and 71.5*s* for ComBat(30.8*s*)+sPLS-DA(40.6*s*).

### 2.4 Study-specific outputs with *MINT*

One of the main challenges when combining independent studies is to assess the concordance between studies. During the integration procedure, MINT proposes not only individual performance accuracy assessment, but also insightful graphical outputs that are study-specific and can serve as Quality Control step to detect outlier studies. One particular example is the Takahashi study from the stem cell data, whose poor performance (Figure 1D) was further confirmed on the study-specific outputs (Figure S9). Of note, this study was the only one generated through Agilent technology and its sample size only accounted for 4.2% of the training set.

The sample plots from each individual breast cancer data set showed the strong ability of MINT to discriminate the breast cancer subtypes while integrating data sets generated from disparate transcriptomics platforms, microarrays and RNA-sequencing (Figure 3A-C). Those data sets were all differently pre-processed, and yet MINT was able to model an overall agreement between all studies; *MINT* successfully built a space based on a handful of genes in which samples from each study are discriminated in a homogenous manner.

**Figure 3:**
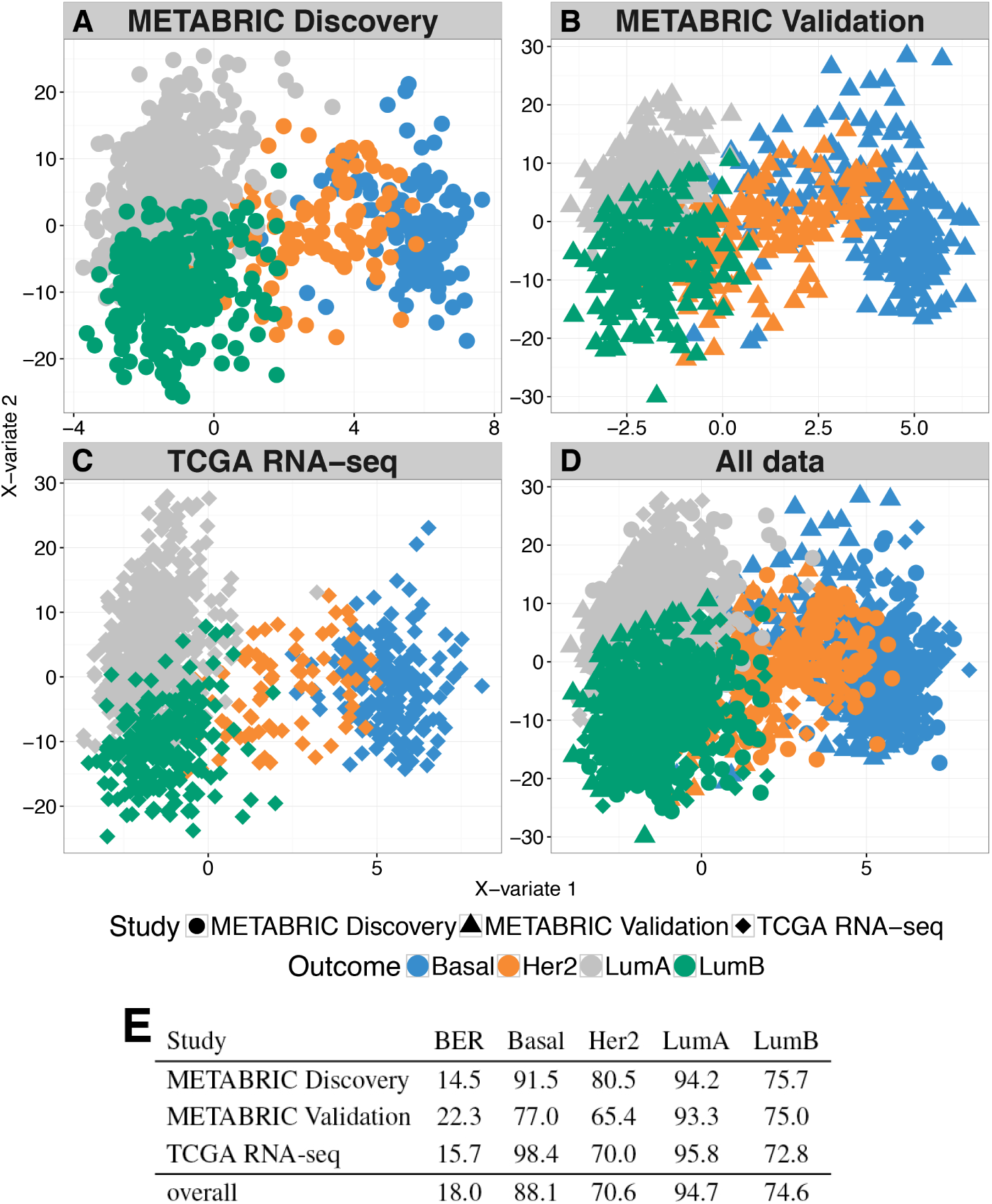
*MINT* study-specific sample plots showing the projection of samples from A) METABRIC Discovery, (B) METABRIC Validation and (C) TCGA-RNA-seq experiments, in the same subspace spanned by the first two MINT components. The same subspace is also used to plot the (D) overall (integrated) data. (E) Balanced Error Rate and classification accuracy for each study and breast cancer subtype from the MINT analysis.

### 2.5 *MINT* gene signature identified promising biomarkers

MINT is a multivariate approach that builds successive components to discriminate all categories (classes) indicated in an outcome variable. On the stem cell data, *MINT* selected 2 and 15 genes on the first two components respectively (Supplemental Information S5.4). The first component clearly segregated the pluripotent cells (fibroblasts) vs. the two non-pluripotent cell types (hiPSC and hESC) (Figure 1C, 1D). Those non pluripotent cells were subsequently separated on component two with some expected overlap given the similarities between hiPSC and hESC. The two genes selected by MINT on component 1 were Lin28A and CAR which were both found relevant in the literature. Indeed, Lin28A was shown to be highly expressed in ESCs compared to Fibroblasts (Yu *et al.*, 2007; Tsialikas and Romer-Seibert, 2015) and CAR has been associated to pluripotency (Krivega *et al.*, 2014). Finally, despite the high heterogeneity of hiPSC cells included in this study, MINT gave a high accuracy for hESC and hiPSC on independent test sets (93.9% and 77.9% respectively, Figure S6), suggesting that the 15 genes selected by MINT on component 2 have a high potential to explain the differences between those cell types (Table S5).

On the breast cancer study, we performed two analyses which either included or discarded the PAM50 genes that were used to define the four cancer subtypes *Basal, HER2, Luminal A* and *Luminal B* (Parker *et al.*, 2009). In the first analysis, we aimed to assess the ability of MINT to specifically identify the PAM50 key driver genes. *MINT* successfully recovered 37 of the 48 PAM50 genes present in the data (77%) on the first three components (7, 20 and 10 respectively). The overall signature included 30, 572 and 636 genes on each component, i.e. 7.8% of the total number of genes in the data. The performance of *MINT* (BER of 17.8 on the training set and 11.6 on the test set) was superior than when performing a PLS-DA on the PAM50 genes only (BER of 20.8 on the training set and a very high 75 on the test set). This result shows that the genes selected by MINT offer a complementary characterisation to the PAM50 genes.

In the second analysis, we aimed to provide an alternative signature to the PAM50 genes by ommitting them from the analysis. *MINT* identified 11, 272 and 253 genes on the first three components respectively (Table S6, S6-2 and Figure S13-S14). The genes selected on the first component gradually differentiated *Basal, HER2* and *Luminal A/B*, while the second component genes further differentiated *Luminal A* from *Luminal B* (Figure 3D). The classification performance was similar in each study (Figure 3E), highlighting an excellent reproducibility of the biomarker signature across cohorts and platforms.

Among the 11 genes selected by MINT on the first component, GATA3 is a transcription factor that regulates luminal epithelial cell differentiation in the mammary glands (Kouros-Mehr *et al.*, 2006; Asselin-Labat *et al.*, 2007), it was found to be implicated in luminal types of breast cancer (Jiang *et al.*, 2014) and was recently investigated for its prognosis significance (McCleskey *et al.*, 2015). The MYB-protein plays an essential role in Haematopoiesis and has been associated to Carcinogenesis (Vargova *et al.*, 2011; Khan *et al.*, 2015). Other genes present in our *MINT* gene signature include XPB1 (Chen *et al.*, 2014), AGR3 (Garczyk *et al.*, 2015), CCDC170 (Yamamoto-Ibusuki *et al.*, 2015) and TFF3 (May and Westley, 2015) that were reported as being associated with breast cancer. The remaining genes have not been widely associated with breast cancer. For instance, TBC1D9 has been described as over expressed in cancer patients (Andres *et al.*, 2013, 2014). DNALI1 was first identified for its role in breast cancer in Parris *et al.* (2010) but there was no report of further investigation. Although AFF3 was never associated to breast cancer, it was recently proposed to play a pivotal role in adrenocortical carcinoma (Lefevre *et al.*, 2015). It is worth noting that these 11 genes were all included in the 30 genes previously selected when the PAM50 genes were included, and are therefore valuable candidates to complement the PAM50 gene signature as well as to further characterise breast cancer subtypes.

## 3 Discussion

There is a growing need in the biological and computational community for tools that can integrate data from different microarray platforms with the aim of classifying samples (integrative classification). Although several efficient methods have been proposed to address the unwanted systematic variation when integrating data (Gagnon-Bartsch and Speed, 2012; Johnson *et al.*, 2007; Sims *et al.*, 2008; Listgarten *et al.*, 2010; Lê Cao *et al.*, 2014), these are usually applied as a pre-processing step before performing classification. Such sequential approach may lead to overfitting and over-optimistic results due to the use of transductive modelling (such as prediction based on ComBat-normalised data (Hughey and Butte, 2015)) and the use of a test set that is normalised or pre-processed with the training set. To address this crucial issue, we proposed a new Multivariate INTegrative method, MINT, that simultaneously corrects for batch effects, classifies samples and selects the most discriminant biomarkers across studies.

MINT seeks to identify a common projection space for all studies that is defined on a small subset of discriminative variables and that display an analogous discrimination of the samples across studies. Therefore, MINT provides sample plot and classification performance specific to each study (Figure 3). Among the compared methods, MINT was found to be the fastest and most accurate method to integrate and classify data from different microarray and RNA-seq platforms. In addition to the integrative classification framework, MINT was extended to an integrative regression framework (multiple multivariate regression, Supplemental Information S2).

Integrative approaches such as MINT are essential when combining multiple studies of complex data to limit spurious conclusions from any downstream analysis. Current methods showed a high proportion of false positives (44% on MAQC data) and exhibited very poor prediction accuracy (PLS-DA, sPLS-DA and RF, Figure 2). For instance, RF was ranked second only to MINT on the breast cancer learning set, but it was ranked as the worst method on the test set. This reflects the absence of controlling for batch effects in these methods and supports the argument that assessing the presence of batch effects is a key preliminary step. Failure to do so, as shown in our study, can result in poor reproducibility of results in subsequent studies, and this would not be detected without an independent test set.

We assessed the ability of *MINT* to identify relevant gene signatures that are reproducible and platform-agnostic. MINT successfully integrated data from the MAQC project by selecting true positives genes that were also differentially expressed in each experiment. We also assessed MINT's capabilities analysing stem cells and breast cancer data. In these studies, MINT displayed the highest classification accuracy in the training sets and the highest prediction accuracy in the testing sets, when compared to sixteen sequential procedures (Figure 2). These results suggest that, in addition to being highly predictive, the discriminant variables identified by MINT are also of strong biological relevance.

In the stem cell data, MINT identified 2 genes LIN28A and CAR, to discriminate pluripotent cells (fibroblasts) against non-pluripotent cells (hiPSC and hESC). Pluripotency is well-documented in the literature and OCT4 is currently the main known marker for undifferentiated cells (Rosner *et al.*, 1990; Schöler *et al.*, 1990; Niwa *et al.*, 2000; Matin *et al.*, 2004). However, MINT did not selected OCT4 on the first component but instead, identified two markers, LIN28A and CAR, that were ranked higher than OCT4 in the DEG list obtained on the concatenated data. While the results from MINT still supported OCT4 as a marker of pluripotency, our analysis suggests that LIN28A and CAR are stronger reproducible markers of differentiated cells, and could therefore be superior as substitutions or complements to OCT4. Experimental validation would be required to further assess the potential of LIN28A or CAR as efficient markers.

Several important issues require consideration when dealing with the general task of integrating data. First and foremost, sample classification is crucial and needs to be well defined. This required addressing in analyses with the stem cell and breast cancer studies generated from multiple research groups and different microarray and RNA-seq platforms. For instance, the breast cancer subtype classification relied on the PAM50 intrinsic classifier proposed by Parker *et al.* (2009), which we admit is still controversial in the literature (Curtis *et al.*, 2012). Similarly, the biological definition of hiPSC differs across research groups (Bilic and Belmonte, 2012; Newman and Cooper, 2010), which results in poor reproducibility among experiments and makes the integration of stem cell studies challenging (Rohart *et al.* 2016). The expertise and exhaustive screening required to homogeneously annotate samples hinders data integration, and because it is a process upstream to the statistical analysis, data integration approaches, including MINT, can not address it.

A second issue in the general process of integrating datasets from different sources is data access and normalisation. As raw data are often not available, this results in integration of data sets that have each been normalised differently, as was the case with the breast cancer data in our study. Despite this limitation, MINT produced satisfactory results in that study. We were also able to overcome this issue in the stem cells data by using the stemformatics resource (Wells *et al.*, 2013) where we had direct access to homogeneously pre-processed data (background correction, log2- and YuGene-transformed Lê Cao *et al.* 2014). In general, variation in the normalisation processes of different data sets produces unwanted variation between studies and we recommend this should be avoided if possible.

A final important issue in data integration involves accounting for both between-study differences and platform effects. When samples clustered by study and the studies clustered by platform, then the experimental platform and not the study, is the biggest source of variation (e.g. 75% of the variance in the breast cancer data, Figure S10A). Indeed, there are inherent differences between commercial platforms that greatly magnify unwanted variability, as was discussed by Shi *et al.* (2006) on the MAQC project. As platform information and study effects are nested, *MINT* and other data integration methods dismiss the platform information and focus on the study effect only. Indeed, each study is considered as included in a single platform. MINT successfully integrated microarray and RNA-seq data, which supports that such an approach will likely be sufficient in most scenarios.

When applying *MINT*, additional considerations need be taken into account. In order to reduce unwanted systematic variation, the method centers and scales each study as an initial step, similarly to BMC (Sims *et al.*, 2008). Therefore, only studies with a sample size > 3 can be included, either in a training or test set. In addition, all outcome categories need to be represented in each study. Indeed, neither MINT nor any classification methods can perform satisfactorily in the extreme case where each study only contains a specific outcome category, as the outcome and the study effect can not be distinguished in this specific case.

## 4 Conclusion

We introduced MINT, a novel Multivariate INTegrative method, that is the first approach to integrate independent transcriptomics studies from different microarray and RNA-seq platforms by *simultaneously*, correcting for batch effects, classifying samples and identifying key discriminant variables. We first validated the ability of MINT to select true positives genes when integrating the MAQC data across different platforms. Then, MINT was compared to sixteen sequential approaches and was shown to be the fastest and most accurate method to discriminate and predict three human cell types (human Fibroblasts, human Embryonic Stem Cells and human induced Pluripotent Stem Cells) and four subtypes of breast cancer (Basal, HER2, Luminal A and Luminal B). The gene signatures identified by MINT contained existing and novel biomarkers that were strong candidates for improved characterisation the phenotype of interest. In conclusion, MINT enables reliable integration and analysis of independent genomic data sets, outperforms existing available sequential methods, and identifies reproducible genetic predictors across data sets. MINT is available through the mixMINT module in the mixOmics R-package.

## 5 Methods

We use the following notations. Let *X* denote a data matrix of size *N* observations (rows) × *P* variables (e.g. gene expression levels, in columns) and *Y* a dummy matrix indicating each sample class membership of size *N* observations (rows) × *K* categories outcome (columns). We assume that the data are partitioned into *M* groups corresponding to each independent study *m*: {(*X*^(1)^, *Y*^(1)^), …, (*X*^(*M*)^, *Y*^(*M*)^)} so that 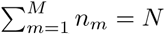, where *n*_*m*_ is the number of samples in group *m*, see Figure 4. Each variable from the data set *X*^(*m*)^ and *Y*^(*m*)^ is centered and has unit variance. We write *X* and *Y* the concatenation of all *X*^(*m*)^ and *Y*^(*m*)^ respectively. For 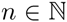, we denote for all 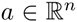 its *l*^1^ norm 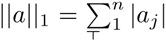 and its *l*^2^ norm 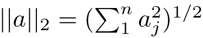 and |a|_+_ the positive part of *a*. For any matrix we denote by its transpose.

**Figure 4:**
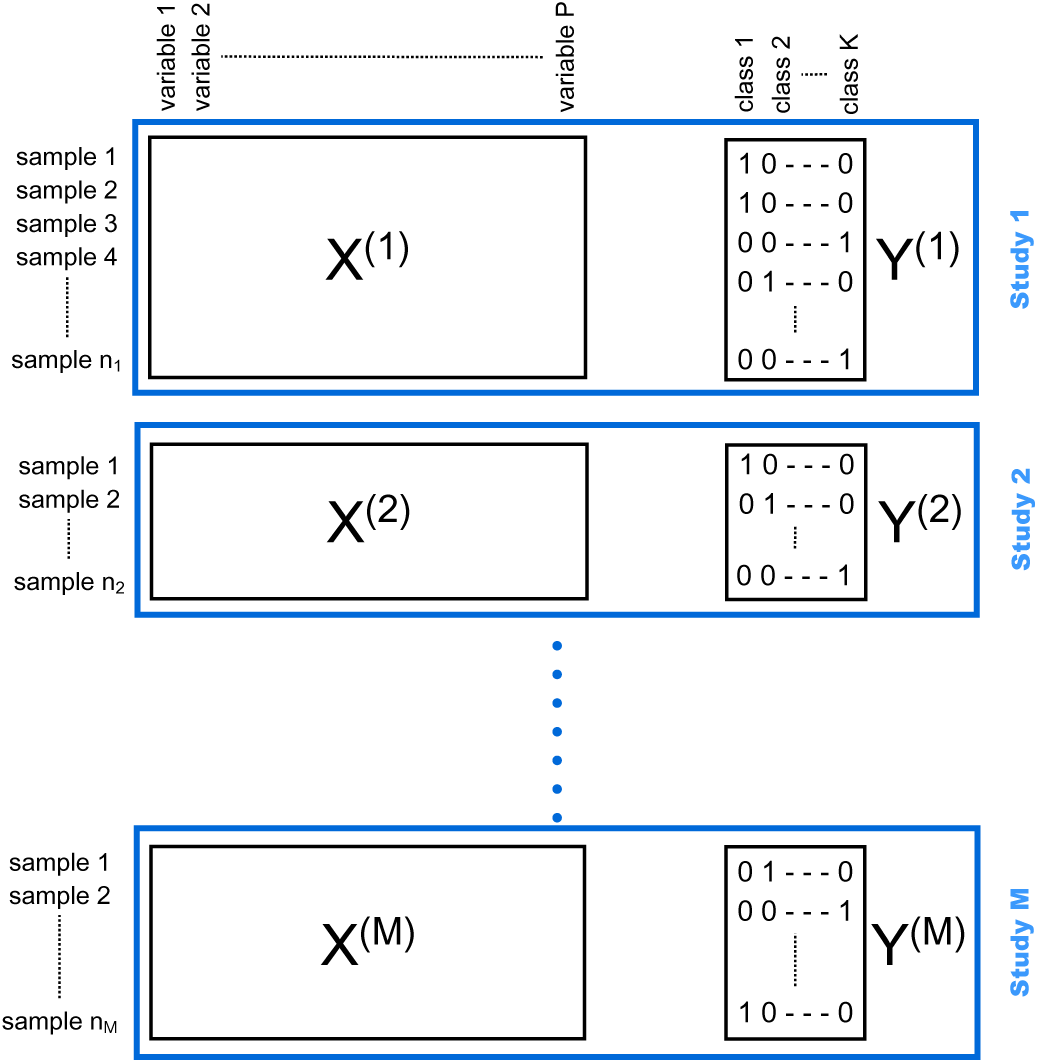
Experimental design of *MINT*, combining *M* independent studies *X*^(*m*)^, *Y*^(*m*)^, where *X*^(*m*)^ is a data matrix of size *n*_*m*_ observations (rows) × *P* variables (e.g. gene expression levels, in columns) and *Y*^(*m*)^ is a dummy matrix indicating each sample class membership of size *n*_*m*_ observations (rows) × *K* categories outcome (columns).

### 5.1 PLS-based classification methods to combine independent studies

PLS approaches have been extended to classify samples *Y* from a data matrix *X* by maximising a formula based on their covariance. Specifically, latent components are built based on the original *X* variables to summarise the information and reduce the dimension of the data while discriminating the Y outcome. Samples are then projected into a smaller space spanned by the latent component. We first detail the classical PLS-DA approach and then describe mgPLS, a PLS-based model we previously developed to model a group (study) structure in *X*.

#### PLS-DA

Partial Least Squares Discriminant Analysis (Barker and Rayens, 2003) is an extension of PLS for a classification frameworks where *Y* is a dummy matrix indicating sample class membership. In our study, we applied PLS-DA as an integrative approach by naively concatenating all studies. Briefly, PLS-DA is an iterative method that constructs *H* successive artificial (latent) components *t*_*h*_ = *X*_*h*_*a*_*h*_ and *u*_*h*_=*Y*_*h*_*b*_*h*_ for *h*=1*, &, H*, where the *h*^*th*^ component *t*_*h*_ (respectively *u*_*h*_) is a linear combination of the *X* (*Y*) variables. *H* denotes the dimension of the PLS-DA model. The weight coefficient vector *a*_*h*_ (*b*_*h*_) is the loading vector that indicates the *importance* of each variable to define the component. For each dimension *h*=1*, …, H* PLS-DA seeks to maximize

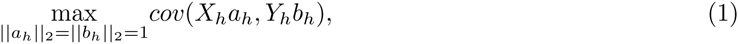

where *X*_*h*_, *Y*_*h*_ are residual matrices (obtained through a *deflation step*, as detailed in Lê Cao *et al.* 2011). The PLS-DA algorithm is described in Supplemental Information S1. The PLS-DA model assigns to each sample *i* a pair of *H* scores 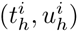 which effectively represents the projection of that sample into the *X*- or *Y* - space spanned by those PLS components. As *H << P*, the projection space is small, allowing for dimension reduction as well as insightful sample plot representation (e.g. graphical outputs in Section 2). While PLS-DA ignores the data group structure inherent to each independent study, it can give satisfactory results when the between groups variance is smaller than the within group variance or when combined with extensive data subsampling to account for systematic variation across platforms (Rohart *et al.*, 2016).

#### mgPLS

Multi-group PLS is an extension of the PLS framework we recently proposed to model grouped data (Eslami *et al.*, 2013, 2014), which is relevant for our particular case where the groups represent independent studies. In mgPLS, the PLS-components of each group are constraint to be built based on the same loading vectors in *X* and *Y*. These *global* loading vectors thus allow the samples from each group or study to be projected in the same common space spanned by the PLS-components. We extended the original unsupervised approach to a supervised approach by using a dummy matrix *Y* as in PLS-DA to classify samples while modelling the group structure. For each dimension *h*=1, …, *H* mgPLS-DA seeks to maximize

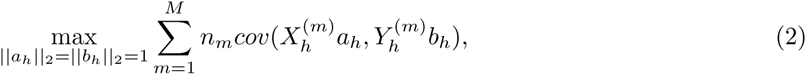

where *a*_*h*_ and *b*_*h*_ are the global loadings vectors common to all groups, 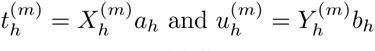 are the group-specific (partial) PLS-components, and 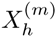 and 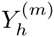 are the residual (deflated) matrices. The global loadings vectors (*a*_*h*_, *b*_*h*_) and global components (*t*_*h*_=*X*_*h*_*a*_*h*_, *u*_*h*_=*Y_h_b_h_*) enable to assess overall classification accuracy, while the group-specific loadings and components provide powerful graphical outputs for each study that is integrated in the analysis. Global and group-specific components and loadings are represented in Figure S1. The next development we describe below is to include internal variable selection in mgPLS-DA for large dimensional data sets.

### 5.2 MINT

Our novel multivariate integrative method *MINT simultaneously* integrates independent studies and selects the most discriminant variables to classify samples and predict the class of new samples. MINT seeks for a common projection space for all studies that is defined on a small subset of discriminative variables and that display an analogous discrimination of the samples across studies. The identified variables share common information across all studies and therefore represent a reproducible signature that helps characterising biological systems. *MINT* further extends mgPLS-DA by including a *l*^1^-penalisation on the global loading vector *a*_*h*_ to perform variable selection. For each dimension *h*=1*, …, H* the *MINT* algorithm seeks to maximize

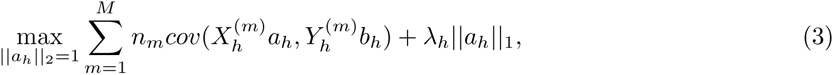

where in addition to the notations from equation (2), *λ_h_* is a non negative parameter that controls the amount of shrinkage on the global loading vectors *a*_*h*_ and thus the number of non zero weights. Similarly to Lasso (Tibshirani, 1996) or sparse PLS-DA (Lê Cao *et al.*, 2011), the added *l*^1^ penalisation in *MINT* improves interpretability of the PLS-components that are now defined only on a set of selected biomarkers from *X* (with non zero weight) that are identified in the linear combination 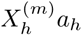. The *l*^1^ penalisation in effectively solved in the *MINT* algorithm using soft-thresholding (see pseudo Algorithm 1).

### 5.3 Class prediction and parameters tuning with *MINT*

MINT centers and scales each study from the training set, so that each variable has mean 0 and variance 1, similarly to any PLS methods. Therefore, a similar pre-processing needs to be applied on test sets. If a test sample belongs to a study that is part of the training set, then we apply the same scaling coefficients as from the training study. If the test study is completely independent, then it is centered and scaled separately.

#### Algorithm 1 MINT

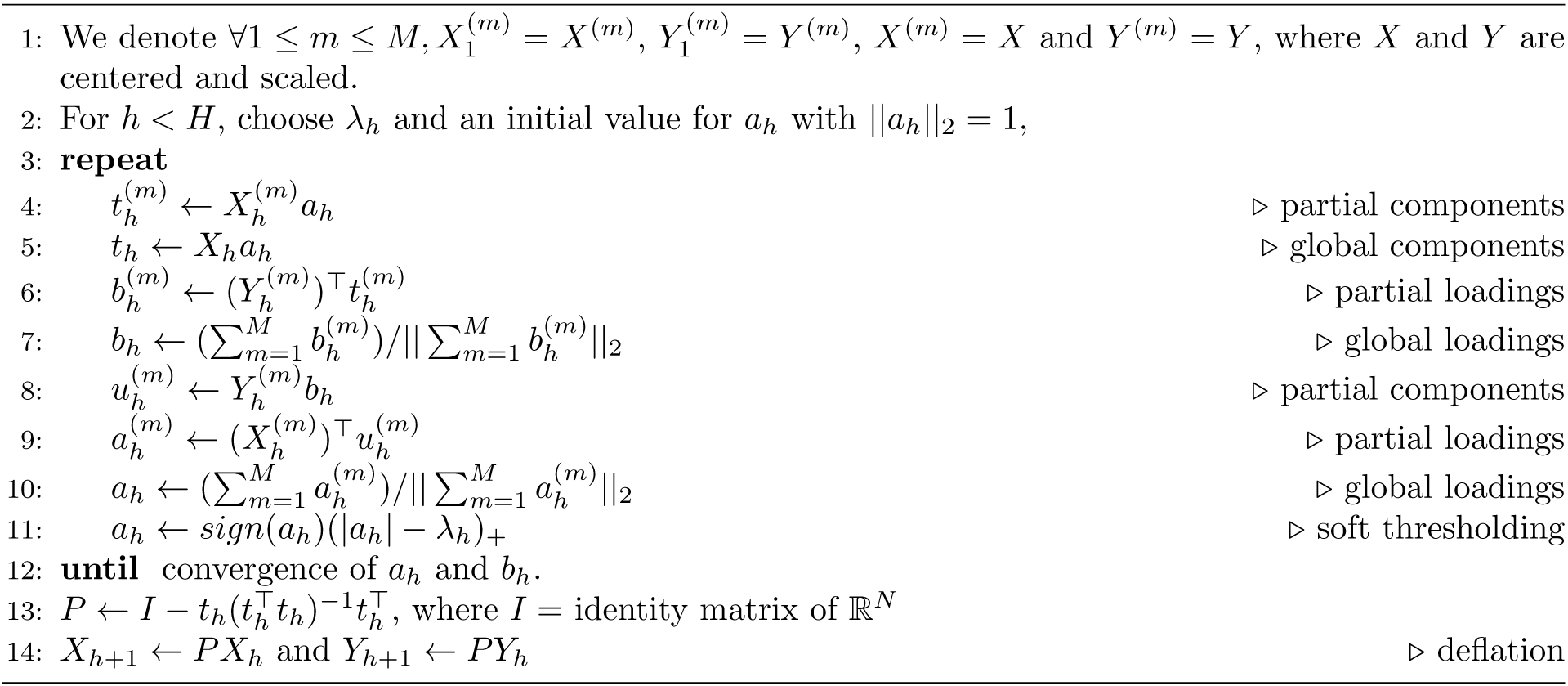

After scaling the test samples, the prediction framework of PLS is used to estimate the dummy matrix *Y*_*test*_ of an independent test set *X*_*test*_ (Tenenhaus, 1998), where each row in *Y*_*test*_ sums to 1, and each column represents a class of the outcome. A class membership is assigned (predicted) to each test sample by using the maximal distance, as described in Lê Cao *et al.* (2011). It consists in assigning the class with maximal positive value in *Y*_*test*_.

The main parameter to tune in MINT is the penalty *λ_h_* for each PLS-component *h*, which is usually performed using Cross-Validation (CV). In practice, the parameter *λ_h_* can be equally replaced by the number of variables to select on each component, which is our preferred user-friendly option. The assessment criterion in the CV can be based on the proportion of misclassified samples, proportion of false or true positives, or, as in our case, the balanced error rate (BER). BER is calculated as the averaged proportion of wrongly classified samples in each class and weights up small sample size classes. We consider BER to be a more objective performance measure than the overall misclassification error rate when dealing with unbalanced classes. *MINT* tuning is computationally efficient as it takes advantage of the group data structure in the integrative study. We used a “Leave-One-Group-Out Cross-Validation (LOGOCV)”, which consists in performing CV where group or study *m* is left out only once *m*=1*, …, M*. LOGOCV realistically reflects the true case scenario where prediction is performed on independent external studies based on a reproducible signature identified on the training set. Finally, the total number of components *H* in *MINT* is set to *K* − 1, *K*=number of classes, similar to PLS-DA and *l*^1^ penalised PLS-DA models (Lê Cao *et al.*, 2011).

### 5.4 Case studies

We demonstrate the ability of *MINT* to identify the true positive genes on the MAQC project, then highlight the strong properties of our method to combine independent data sets in order to identify reproducible and predictive gene signatures on two other biological studies.

#### The MicroArray Quality Control (MAQC) project

The extensive MAQC project focused on assessing microarray technologies reproducibility in a controlled environment (Shi *et al.*, 2006). Two reference samples, RNA samples Universal Human Reference (UHR) and Human Brain Reference (HBR) and two mixtures of the original samples were considered. Technical replicates were obtained from three different array platforms-Illumina, AffyHuGene and AffyPrime- for each of the four biological samples A (100% UHR), B (100% HBR), C (75% UHR, 25% HBR) and D (25%UHR and 75% HBR). Data were downloaded from Gene Expression Omnibus (GEO) - GSE56457. In this study, we focused on identifying biomarkers that discriminate A vs. B and C vs. D. The experimental design is referenced in Table S1.

#### Stem cells

We integrated 15 transcriptomics microarray datasets to classify three types of human cells: human Fibroblasts (Fib), human Embryonic Stem Cells (hESC) and human induced Pluripotent Stem Cells (hiPSC). As there exists a biological hierarchy among these three cell types, two sub-classification problems are of interest in our analysis, which we will address simultaneously with *MINT*. On the one hand, differences between pluripotent (hiPSC and hESC) and non-pluripotent cells (Fib) are well-characterised and are expected to contribute to the main biological variation. Our first level of analysis will therefore benchmark *MINT* against the gold standard in the field. On the other hand, hiPSC are genetically reprogrammed to behave like hESC and both cell types are commonly assumed to be alike. However, differences have been reported in the literature (Bilic and Belmonte, 2012; Chin *et al.*, 2009; Newman and Cooper, 2010), justifying the second and more challenging level of classification analysis between hiPSC and hESC. We used the cell type annotations of the 342 samples as provided by the authors of the 15 studies. The stem cell dataset provides an excellent showcase study to benchmark *MINT* against existing statistical methods to solve a rather ambitious classification problem.

Each of the 15 studies was assigned to either a training or test set. Platforms uniquely represented were assigned to the training set and studies with only one sample in one class were assigned to the test set. Remaining studies were randomly assigned to training or test set. Eventually, the training set included eight datasets (210 samples) derived on five commercial platforms and the independent test set included the remaining seven datasets (132 samples) derived on three platforms (Table 1 and Table S3).

**Table 1:** Stem cell experimental design. A total of 15 studies were analysed, including three human cell types, human Fibroblasts (Fib), human Embryonic Stem Cells (hESC) and human induced Pluripotent Stem Cells (hiPSC) across five different types of microarray platforms. Eight studies from five microarray platforms were considered as a training set and seven independent studies from three of the five platforms were considered as a test set.

| Experiment | Platform | Fib | hESC | hiPSC |
| --- | --- | --- | --- | --- |
| Bock | Affymetrix HT-HG-U133A | 6 | 20 | 12 |
| Briggs | Illumina HumanHT-12 V4 | 18 | 3 | 30 |
| Chung | Affymetrix HuGene-1.0-ST V1 | 3 | 8 | 10 |
| Ebert | Affymetrix HG-U133 Plus2 | 2 | 5 | 3 |
| Guenther | Affymetrix HG-U133 Plus2 | 2 | 17 | 20 |
| Maherali | Affymetrix HG-U133 Plus2 | 3 | 3 | 15 |
| Marchetto | Affymetrix HuGene-1.0-ST V1 | 6 | 3 | 12 |
| Takahashi | Agilent SurePrint G3 GE 8x60K | 3 | 3 | 3 |
| Total training set | 5 platforms | 43 | 62 | 105 |
| Andrade | Affymetrix HuGene-1.0-ST V1 | 3 | 6 | 15 |
| Hu | Affymetrix HG-U133 Plus2 | 1 | 5 | 12 |
| Kim | Affymetrix HG-U133 Plus2 | 1 | 1 | 3 |
| Loewer | Affymetrix HG-U133 Plus2 | 4 | 2 | 7 |
| Si-Tayeb | Affymetrix HG-U133 Plus2 | 3 | 6 | 6 |
| Vitale | Illumina HumanHT-12 V4 | 8 | 3 | 18 |
| Yu | Affymetrix HG-U133 Plus2 | 2 | 10 | 16 |
| Total test set | 3 platforms | 22 | 33 | 77 |

The pre-processed files were downloaded from the www.stemformatics.org collaborative platform (Wells *et al.*, 2013). Each dataset was background corrected, log2 transformed, YuGene normalized and mapped from probes ID to Ensembl ID as previously described in Lê Cao *et al.* (2014), resulting in 13 313 unique Ensembl gene identifiers. In the case where datasets contained multiple probes for the same Ensembl ID gene, the highest expressed probe was chosen as the representative of that gene in that dataset. The choice of YuGene normalisation was motivated by the need to normalise each sample independently rather than as a part of a whole study (e.g. existing methods ComBat (Johnson *et al.*, 2007), quantile normalisation (RMA, Bolstad *et al.* 2003)), to effectively limit over-fitting during the CV evaluation process.

#### Breast Cancer

We combined whole-genome gene-expression data from two cohorts from the Molecular Taxonomy of Breast Cancer International Consortium project (METABRIC, Curtis *et al.* (2012) and of two cohorts from the Cancer Genome Atlas (TCGA, Cancer Genome Atlas Network and others (2012)) to classify the intrinsic subtypes *Basal, HER2, Luminal A* and *Luminal B*, as defined by the PAM50 signature (Parker *et al.*, 2009). The METABRIC cohorts data were made available upon request, and were processed by Curtis *et al.* (2012). TCGA cohorts are gene-expression data from RNA-seq and microarray platforms. RNA-seq data were normalised using Expectation Maximisation (RSEM) and percentile-ranked gene-level transcription estimates. The microarray data were processed as described in Cancer Genome Atlas Network and others (2012).

The training set consisted in three cohorts (TCGA RNA-seq and both METABRIC microarray studies), including the expression levels of 15 803 genes on 2 814 samples; the test set included the TCGA microarray cohort with 254 samples (Table 2) Two analyses were conducted, which either included or discarded the PAM50 genes from the data. The first analysis aimed at recovering the PAM50 genes used to classify the samples. The second analysis was performed on 15, 755 genes and aimed at identifying an alternative signature to the PAM50.

**Table 2:** Experimental design of four breast cancer cohorts including 4 cancer subtypes: *Basal, HER2, Luminal A* (LumA) and *Luminal B* (LumB).

| Experiment | Platform | Basal | Her2 | LumA | LumB |
| --- | --- | --- | --- | --- | --- |
| METABRIC Discovery | Illumina HT-12 v3 | 118 | 87 | 466 | 268 |
| METABRIC Validation | Illumina HT-12 v3 | 213 | 153 | 255 | 224 |
| TCGA RNA-seq | illumina HiSeq 2000 | 188 | 80 | 549 | 213 |
| Total training set | 2 platforms | 519 | 320 | 1270 | 705 |
| TCGA microarray | Agilent custom 244K | 57 | 31 | 99 | 67 |
| Total test set | 1 platform | 57 | 31 | 99 | 67 |

### 5.5 Performance comparison with sequential classification approaches

We compared *MINT* with sequential approaches that combine batch-effect removal approaches with classification methods. As a reference, classification methods were also used on their own on a naive concatenation of all studies. Batch-effect removal methods included Batch Mean-Centering (BMC, Sims *et al.* 2008), ComBat (Johnson *et al.*, 2007), linear models (LM) or linear mixed models (LMM), and classification methods included PLS-DA, sPLS-DA (Lê Cao *et al.*, 2011), mgPLS (Eslami *et al.*, 2013, 2014) and Random forests (RF, Breiman 2001). For LM and LMM, linear models were fitted on each gene and the residuals were extracted as a batch-corrected gene expression (Whitcomb *et al.*, 2010; Rohart *et al.*, 2014). The study effect was set as a fixed effect with LM or as a random effect with LMM. No sample outcome (e.g. cell-type) was included. Prediction with ComBat normalised data were obtained as described in Hughey and Butte (2015). In this study, we did not include methods that require extra information-as control genes with RUV-2 (Gagnon-Bartsch and Speed, 2012)- and methods that are not widely available to the community as LMM-EH (Listgarten *et al.*, 2010). Classification methods were chosen so as to simultaneously discriminate all classes. With the exception of sPLS-DA, none of those methods perform internal variable selection. The multivariate methods PLS-DA, mgPLS and sPLS-DA were run on *K* − 1 components, sPLS-DA was tuned using 5-fold CV on each component. All classification methods were combined with batch-removal method with the exception of mgPLS that already includes a study structure in the model.

MINT and PLS-DA-like approaches use a prediction threshold based on distances (see Section 5.3) that optimally determines class membership of test samples, and as such do not require receiver operating characteristic (ROC) curves and area under the curve (AUC) performance measures. In addition, those measures are limited to binary classification which do not apply for our stem cell and breast cancer multi-class studies. Instead we use Balanced classification Error Rate to objectively evaluate the classification and prediction performance of the methods for unbalanced sample size classes (Section 5.2). Classification accuracies for each class were also reported.

## Acknowledgements

This project was partly funded by the ARC Discovery grant project DP130100777 and the Australian Cancer Research Foundation for the Diamantina Individualised Oncology Care Centre at UQDI (FR), and the National Health and Medical Research Council (NHMRC) Career Development fellowship (APP1087415) (KALC).

